# The Connectivity Index: An Effective Metric for Grading Epileptogenicity

**DOI:** 10.1101/635664

**Authors:** Qi Yan, Nicolas Gaspard, Hitten P Zaveri, Hal Blumenfeld, Lawrence J. Hirsch, Dennis D. Spencer, Rafeed Alkawadri

## Abstract

**Objective:** To investigate the performance of a metric of functional connectivity to classify and grade the excitability of brain regions based on evoked potentials to single pulse electrical stimulation (SPES).

**Methods:** Patients who received 1-Hz frequency stimulation between 2003 and 2014 at Yale at prospectively selected contacts were included. The stimulated contacts were classified as seizure onset zone (SOZ), highly irritative zone (IZ_p_) or control. Response contacts were classified as seizure onset zone (SOZ), active interictal (IZ_p_), quiet or other. The normalized number of responses was defined as the number of contacts with any evoked responses divided by the total number of recorded contacts, and the normalized distance is the ratio of the average distance between the site of stimulation and sites of evoked responses to the average distances between the site of stimulation and all other recording contacts. A new metric we labeled the connectivity index (CI) is defined as the product of the two values.

**Results:** 57 stimulation-sessions in 22-patients were analyzed. The connectivity index (CI) of the SOZ was higher than control (median CI of 0.74 vs. 0.16, p = 0.0002). The evoked responses after stimulation of SOZ were seen at further distance compared to control (median normalized distance 0.96 vs. 0.62, p = 0.0005). It was 1.8 times more likely to record a response at SOZ than in non-epileptic contacts after stimulation of a control site. Habitual seizures were triggered in 27% of patients and 35 % of SOZ contacts (median stimulation intensity 4 mA) but in none of the control or IZ_p_ contacts. Non-SOZ contacts in multifocal or poor surgical outcome cases had a higher CI than non-SOZ contacts in those with localizable onsets (medians CI of 0.5 vs. 0.12, p = 0.04). There was a correlation between the stimulation current intensity and the normalized number of evoked responses (r = + 0.49, p 0.01) but not with distance (r = + 0.1, p 0.64)

**Conclusions:** We found enhanced connectivity when stimulating the SOZ compared to stimulating control contacts; responses were more distant as well. Habitual auras and seizures provoked by SPES were highly predictive of brain sites involved in seizure generation.

## Introduction

Victor Horsley used faradic electrical stimulation to confirm the localization of the epileptic focus in one of John Hughlings Jackson’s patient who underwent resection of an epileptic focus in 1886^18^. Harvey Cushing used this technique in 1909 to define the sensorimotor cortex surrounding a tumor and to confirm the localization of epileptic seizures that manifested with sensory auras^5^ and predicted that stimulation will be invaluable for localization of epileptic foci. Presently, recording of seizures and localization of the seizure onset zone (SOZ) in the subset of surgical patients who require intracranial EEG (icEEG) evaluations remains the best available means to identify epileptogenic tissue^3^. Intracranial EEG presents two significant challenges: i) It is invasive and may have serious complications, which increase in rate as a function of the duration of monitoring^6^, and ii) The chances of sustained seizure freedom after epilepsy surgery falls to between 30-80% depending on the lobe involved. Hence, studies that enhance on effective localization in the interictal phase are still needed. These challenges suggest that current methods of localization are not optimal and that defining epileptogenicity based on seizure-onset zones, while practical, falls short of ‘ideal’^7,9,15^. Given the aforementioned limitations and associated risks in current standards, identifying reliable markers of epileptogenicity is desired.

Based on known mechanisms of imbalance between excitatory and inhibitory responses especially at the level of abnormal synaptic plasticity that could transform high-normal levels of network activity into ictal activity and supported by our direct bedside observation in-vivo. And based on known mechanisms of generation of EEG/evoked potentials and the post-synaptic level and recent findings in literature - We aimed to test the performance of a novel metric to gauge the regional and global facilitation of synaptic transmission and hence in theory the overall epileptogenic potential based on evoked the normalized number of evoked responses to single pulse stimulation weighted by the distance it was recorded at to minimize the effect of regional sampling inhomogeneity.

A few studies have demonstrated that Single Pulse Electrical Stimulation (SPES) of the epileptogenic region, or its vicinity, results in responses with accentuated amplitudes^8,16,17^. To our knowledge, the spatial extent of responses was not systematically quantified, and correlation with surgical outcomes was not always studied. One issue arises from the limited spatial resolution and sampling of the brain via current methods of icEEG, which varies by the clinical hypothesis and by local epilepsy surgery team preferences.

In this single-center case-control study, we test the relationship between (CI) that is based on the quantity and spatial extent of evoked responses to SPES, and a qualitative classification of contacts. We also estimate the predictive value of focal seizures and auras induced by the low-frequency stimulation (1 Hz) used during SPES in this cohort of drug-resistant epilepsies.

## Methods

### Patients and intracranial EEG data acquisition

We retrospectively identified patients who underwent 1-Hz stimulation at prospectively selected sites for clinical or research purposes at the Yale Comprehensive Epilepsy Center between 2003 and 2014. EEG data were acquired with either Biologic (sampling rate 256-512 Hz) or Natus (1024 Hz) acquisition systems [both now Natus Medical Incorporated, USA]. We reviewed the pre-surgical work up data and notes from stimulation with documentation of clinical events such as side effects, auras or seizures, as well as pathologies and surgical outcomes when available. The recordings and averaging of evoked responses were performed using a referential montage with a reference electrode placed typically in the diploic space. The electrodes used were commercially available platinum-iridium (Ad-Tech, WI) with 2.8 mm exposed surface and 10 mm center to center inter-electrode contact spacing arranged in subdural grids or strips and depth electrodes with 1.1 mm diameter and 10 mm center to center inter-electrode distance placed as needed to target deep structures. Surgical outcomes were classified as ‘good’ (Engel class I and II) and poor (Engel III, IV). We analyzed surgical outcomes at 24 months or after the epilepsy surgery. The coregistration of the contacts with pre-operative MRI space was based on pre-implantation and post-implantation MRIs and post-implantation CT scans using BioImage Suite (v3.0, CT, USA, www.bioimagesuite.org)^10^. Each contact was assigned a lobe (frontal, parietal, temporal, occipital, and insular) and a sub-lobe (lateral frontal LF, medial frontal/cingulate MF/CG, orbitofrontal OF, medial parietal MP, lateral parietal LP, lateral temporal LT, mesial temporal MT (including hippocampus, entorhinal cortex, parahippocampal gyrus), basal temporal BT, lateral occipital LO, medial occipital MO, basal occipital BO) based on its spatial relationship to anatomical landmarks. Each electrode was identified in the MRI-coordinate system with three values (x, y, z). The Euclidian distance between electrodes was used in this study. The implantation usually involved a grid covering the lateral frontal-temporal-parietal convexity, several strips spanning most of the major sub-lobes above, and depth electrodes as needed to target mesial structures.

### Classification of the electrode contacts

The stimulated contacts were classified as: i. seizure onset (SOZ) which was determined by the electrodes that showed the first electroencephalographic ictal changes, ii. within epileptic network/ possibly epileptogenic active irritative zone (IZ_p_, frequent or periodic spiking > 10 discharges per minute during NREM sleep, and early ictal propagation within 5 seconds of seizure onset) or iii. control. The control stimulation contacts were not involved in ictal or interictal epileptic activity and were located > 5 cm in distance from SOZ and IZ contacts. Response contacts were classified as seizure onset zone (SOZ), active interictal (IZ and areas of less frequent interictal epileptiform discharges), quiet or other if information missing to classify. For presentation of the results below, *epileptic contacts* refer to both the SOZ and IZ_p_ contacts, and *non-SOZ* contacts refer to all contacts with the exception of SOZ contacts.

### Single-Pulse Evoked Potentials (SPEPs) and Connectivity Index

The stimulation was performed typically at the end of video-EEG evaluation after seizures were recorded and anti-seizure medications were reinstated while patients are in a relaxed wakeful state. Evoked potentials were estimated by stimulation of a pair of adjacent contacts (i.e. biphasic bipolar stimulation) and recording averaged responses from all remaining contacts (typically between 110 to 254 contacts in data studied here). Current-controlled Grass Technologies S88 or Nicolet Cortical Stimulator (Natus, USA) were used. The stimulation was performed up to 2 minutes at 1 Hz with alternating bipolar 250-300 microsecond square wave pulses, and 0.5-1 mA increments up to 9-12 mA, or lower currents if symptoms or after discharges occurred. EEG signal was band-passed between 1-500 Hz (1-120 Hz if sampling frequency 512 Hz or less) and evoked responses to single pulse stimuli were averaged twice for reproducibility: An evoked response was defined to be reproducible when the correlation coefficient was > 0.9 between two averages. The peaks of the waveforms were marked manually by an expert reader. If present, we calculated the following values for each response according to known characteristics of the cortical evoked responses^12^: First negative peak (N1) latency, second negative peak (N2) latency, inter-peak N1-N2 interval, maximum peak-to-peak amplitude. In complex responses most encountered during stimulation of epileptic sites, we used first and second negative waveforms to conduct this step. We also calculated an index that weighs the normalized number of evoked responses and the normalized distance between the stimulated contacts and the contacts where the responses were recorded at as follows

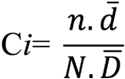

Where (CI) is the Connectivity index, n, the number of contacts with evoked responses, N the total number of contacts, d the average Euclidian distance of contacts with evoked responses from the site of stimulation and D the average Euclidian distance of all contacts from site of stimulation. Distance is factored to emphasize remote responses and to compensate partially for oversampling from SOZ. In the results section normalized number of responses represents n/N, whereas 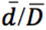, represents normalized distance. If seizures were induced, we employed in averaging segments prior to electrographic onset if any. See figure-1 for schematic illustration of calculating Ci.

**Figure 1:**
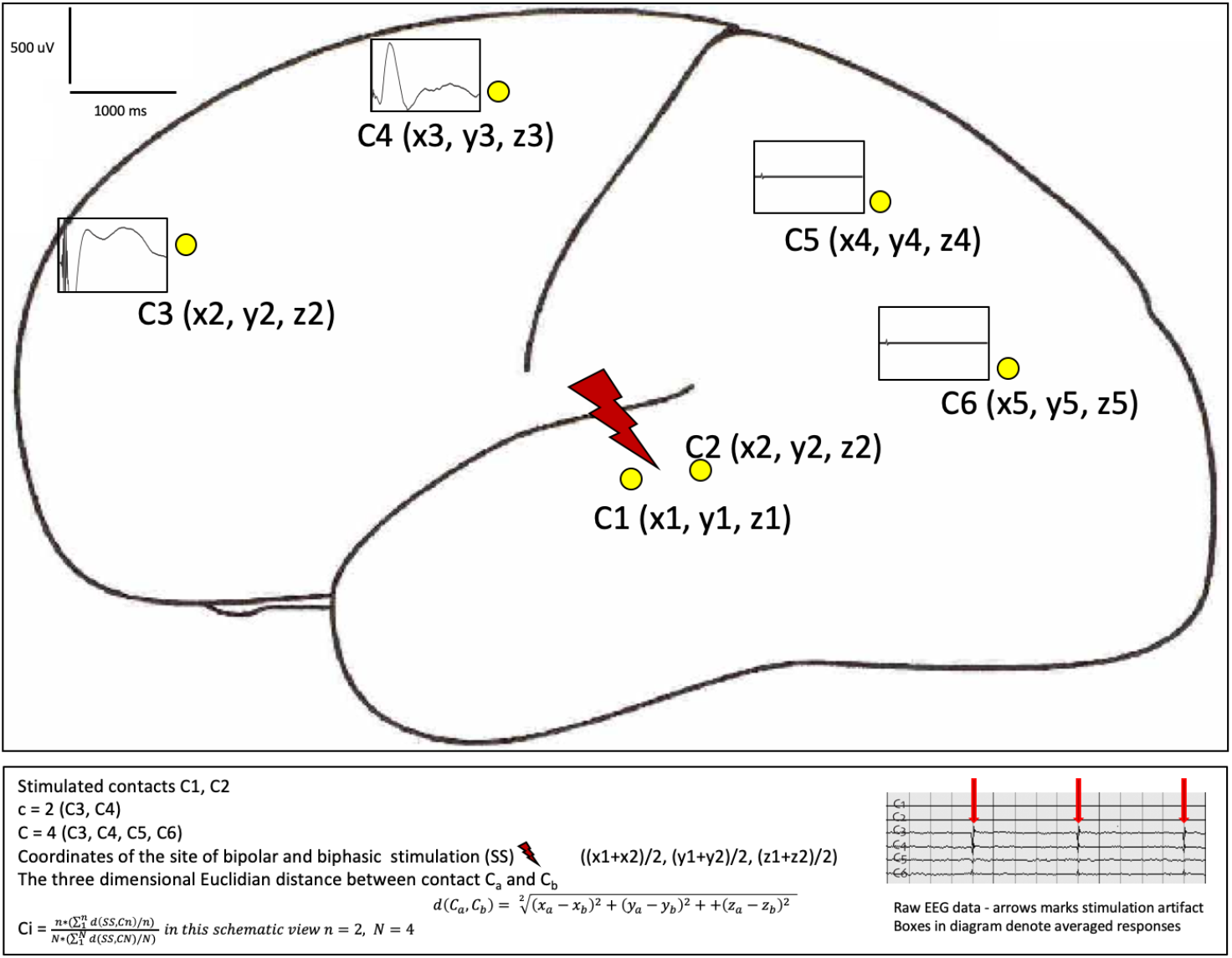
Schematic representation of CI calculation.

For <u>qualitative analysis</u>: Recording channels were classified according to presence or absence of delayed responses defined as evoked responses with peak latencies > 150 ms and duration of 225 ms or greater. The spatial distribution was determined per the following: Focal if seen in 9 adjacent and consecutive contacts or less, regional in 9-20 adjacent areas, multi-regional if observed in > 20 but less than 50 % of contacts, and wide-spread if seen in > 50 % of recording contacts. The study was approved by Yale institutional board review HIC# 1408014451

### Statistical analysis

The Statistical evaluations were implemented in JMP (version 10, SAS Institute Inc. Cary, NC 1998-2010). Fisher’s exact test was used for qualitative contingency tables and odds ratio statistical significance. Receiver operating characteristic (ROC) curves were calculated to estimate the effective cut-off for CI as a classifier. Our accepted pre-test probability of alpha error (p-value) was < 0.05 corresponding as well with 95% confidence intervals.

## Results

57 stimulation sessions (2-5 pairs/session) in 22 patients were analyzed. Age of onset of epilepsy was 0.25 – 44 years. Median age at the time of implantation 32.5 years, and interquartile range (28-42.25). All but one was right-handed (handedness data unavailable in one patient). Evoked responses were recorded from 3019 electrode contacts (mean 137.2/case, SD 52.16, SOZ 127, IZ 362, non-epileptic control 1084, and other/unclassified 1446 contacts). The stimulated contacts were 23 SOZ (1 MF, 1 OF, 2 LP, 12 MT, 2 BT, 5 LT), 13 IZ (3 LF, 1 OF, 1 LP, 2 LT, 3 MT, 2 BT, 1 MO), and 21 controls (3 LF, 3 OF, 1 LP, 1 MP, 2 LT, 1 MT, 1 LO, 7 MO, 2 CG). Fourteen patients underwent resection. Please refer to Table 1 for clinical details.

**Table 1.** summary of demographics, seizure localization, surgical resection and outcomes

numbers in brackets in the second column represent the duration of epilepsy at the time of implantation, all ages are in years. R = right, L = left , N = no, Y = yes, AEDs = Anti-seizure medications , MTS = mesial temporal sclerosis, FP = fronto-polar, FCD: focal cortical dysplasia,
| Age at onset | Age at stimulation | Duration of epilepsy | Frequency SZ/month | Generalized motor seizures | Resection | Outcome** | Pathology |
| --- | --- | --- | --- | --- | --- | --- | --- |
| 0.5 | 33 | 32 | 4 | N | R temporal lobectomy + fusiform | I | MTS |
| 29 | 36 | 7 | 30 | Y (rare) | R temporal Heschel's + tip | III | non-specific gliosis |
| 44 | 53 | 9 | 6 | Y (occasional) | - | NR: high risk memory | - |
| 15 | 33 | 18 | 75 | Y | R Parietal | I | FCD |
| 4 | 36 | 32 | 1.5 | Y | - | NR: > one focus | - |
| 19 | 54 | 34 | 16 | Y (1 in lifetime) | - | NR: > one focus | - |
| 3 | 30 | 27 | 12 | Y (2 in lifetime) | L anterior temporal | Ib | Ganglioglioma + MTS |
| 38 | 47 | 9 | 8 | N | L anterior temporal | III | non-specific |
| 8 | 22 | 14 | 2 | N | L frontal | I | FCD |
| 17 | 28 | 11 | 75 | Y | L temporo-parietal/frontal resection | III-IV | non-specific |
| 19 | 27 | 8 | 8 | Y (nocturnal) | R anterior temporal | III-IV | Neuronal loss HC rare heterotopia |
| 25 | 43 | 18 | 30 | Y | - | NR: > one focus | - |
| 17/0.25 | 31 | 14/31 | 1.5 | Y | - | NR: Non-localizable | - |
| 0.5 | 40 | 39 | 7 | Y | R posterior temporal, inferior parietal | Id | remote ischemic |
| 11/0.5 | 26 | 15/26 | 360 | Y | R pre-cuneus - and MTS FP | III-IV | heterotropic neurons |
| 21 | 28 | 7 | 3 | Y | R lateral TP and post mesial | III-IV | Heterotopia/FCD |
| 2 | 19 | 17 | 3 | N | - | NR: High risk memory | - |
| 9/2 | 17 | 8/15 | 16 | Y | R anterior temporal | III | Limited data |
| 15 | 28 | 13 | 30 | N | - | NR: Non-localizable | - |
| 36 | 53 | 17 | 8 | Y | R frontal | I | Remote ischemic |
| 17 | 32 | 15 | 2 | Y | - | NR: > one focus | - |
| 17 | 48 | 31 | 6 | N | R basal and mesial temporal | I | Gliosis |

\*\*Outcome: Latin numerals in Outcome are seizure outcomes according to Engel.
NR = no resection, X = times, HC = hippocampus. Non-localizable: in reference to onsets involving > 3 sub-lobes simultaneously.

### Connectivity Index (CI) and Distance from site of stimulation

Stimulation of SOZ evoked reproducible responses at significantly higher rates than stimulation of control sites (medians of the normalized number of contacts with evoked responses = n/N, 0.72 vs. 0.26, p 0.0003). These differences were even stronger after normalization to average distance recorded (i.e. the connectivity index CI) (0.74 vs. 0.16, p 0.0002). This holds true when excluding contacts where stimulation led to seizures (medians 0.78 vs. 0.16 p 0.00026). The evoked responses after stimulation of the SOZ contacts were seen at a further distance from the site of stimulation (medians of normalized distances 0.96 vs. 0.62, p 0.0005, median absolute values: 58 mm vs. 48 mm). Medians of CI and normalized distance values for SOZ compared to IZ, however, were not statistically significant. If we excluded cases where stimulation was limited (for example by provoking auras with currents < 5 mA), CI was non-significantly higher in the SOZ contact vs. IZ contacts (0.78 vs. 0.4 p=0.2).

### Connectivity index as a classifier

CI(SOZ) > CI(IZ_p_), and CI(SOZ) > CI(Control) was true in all cases (i.e. within subject) whenever available. The use of CI < 0.19 was 0.92 specific and 0.71 sensitive to classify normal control sites, and CI >0.3 was 0.92 sensitive and 0.70 specific for epileptic contacts.

Qualitative analysis Late and slow evoked responses were almost always seen in epileptic contacts (i.e., SOZ or IZ_p_) (Odds ratio 150, 95% confidence interval 14.4 to 1562). In fact, late and slow responses were observed after stimulation in control sites in only two cases, 1 of which had wide-spread epileptogenicity and active interictal discharges, so no resection took place. Widespread responses were more likely to be recorded from SOZ vs. non-SOZ (odds ratio 41.3, 95% confidence limit 4.3 to 405.4) and SOZ vs. IZ (odds ratio 38, 95% confidence limit, 3.4 to 375.3). See figure 2 for details.

**Figure 2.**
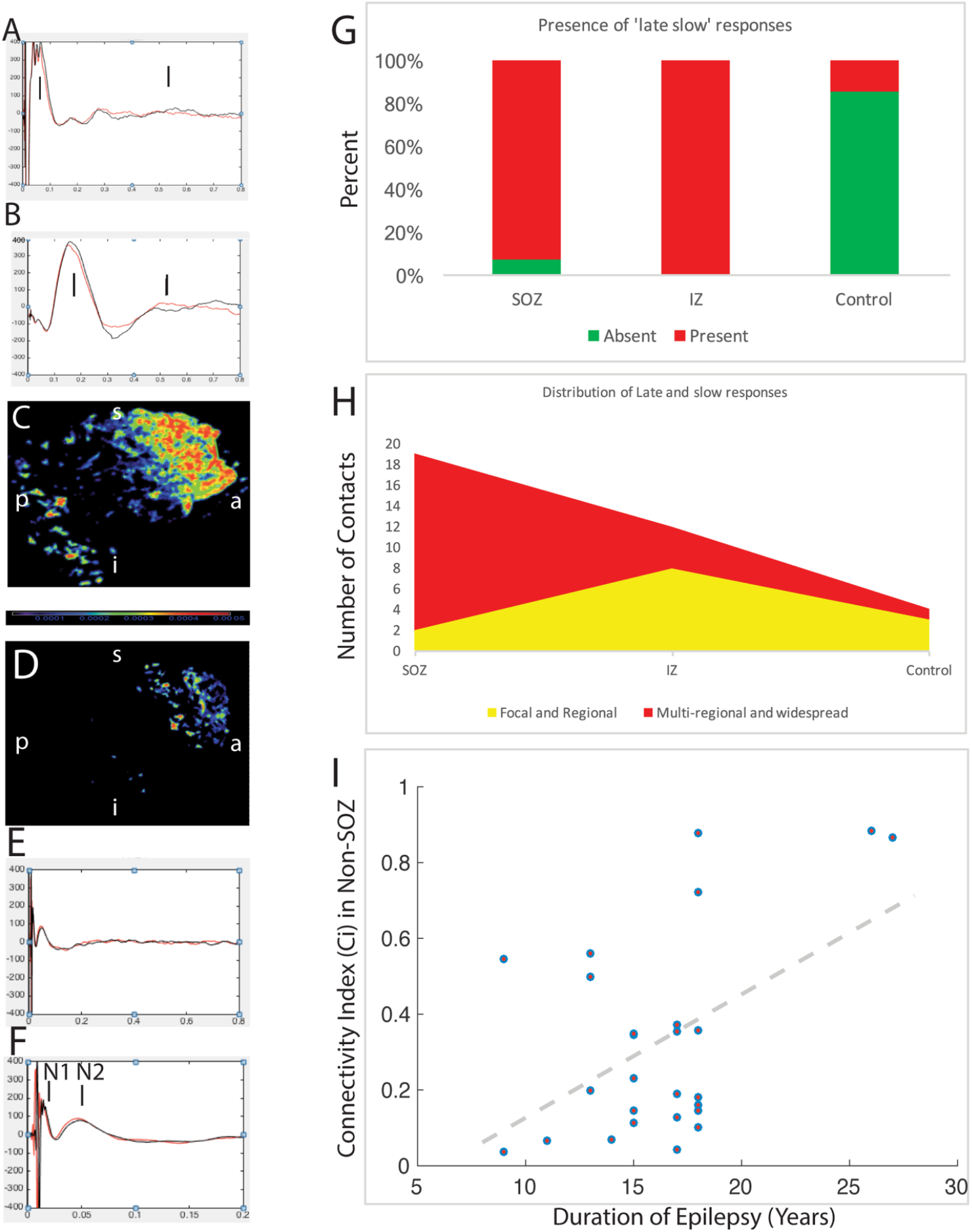
Example of evoked responses from stimulation of epileptic (A, B, C), and stimulation of control non-epileptic single pulse stimulation of the orbitofrontal (D, E, F) regions from two different cases. In A, and B the x axis is 800 milliseconds, and y axis (800 microvolts). Notice the high amplitude and late responses marked by vertical lines. E, and F depict normal physiologic N1, N2 following stimulation of non-epileptic control site on an 800 and 200 millisecond axial scales respectively and 800-microvolts vertical scale. Note that no late responses were recorded. The voltage values are reversed from original signal for a conventional view of EEG (i.e. up is negative). Unlike the predictable N1-N2 configuration in healthy control, stimulation and recording from epileptic contacts results in multiple peaks and is less predictable. To illustrate a volume of normal of pathologic evoked responses (i.e. slow and late), C and D depict volumes based on minimum-norm estimate and source localization of delta waveforms 1-4 Hz between 100-1000 milliseconds after stimulation. The unit of color scale is m.A per m2. a, s, p, i represent anterior, superior, posterior and inferior orientations respectively. G. Ratio of contacts with late and slow evoked responses per contact group H. Qualitative classification of the spatial extent of averaged evoked responses of focal/regional vs. multi-regional or widespread. I. Correlation between duration of epilepsy and CI of non-SOZ contacts; notice the statistically significant correlation r = 0.49 p 0.01.

### Latencies and Duration of Abnormal Evoked Responses and Relation to SOZ

The average latency of N2 (188 ms vs. 167 ms) and duration of N1 (97 ms vs. 81 ms) were significantly longer after stimulation of epileptic contacts compared to control stimulations. There was no difference in latency or amplitude of N1 between the epileptic contacts compared to control (p 0.25 for latency, and 0.29 for amplitude).

It was 1.8 times more likely to record an evoked response from the SOZ than other contacts after stimulation of a control site (p=0.002). It was 2.2 times more likely to observe an N2 response with peak latency > 400 ms after stimulation of an epileptic contact (p=0.0015). Only 5.6 % of N2 responses with latencies > 400 ms were recorded from a quiet site after stimulation of control site. There was a correlation between the current intensity and normalized number of evoked responses (r = + 0.49, p 0.01) but not between current intensity and normalized distance (r = + 0.1, p 0.64).

### Seizures Provoked by Low Frequency Stimulation

Habitual partial seizures or auras were triggered in 27 % of the patients and 35 % of all seizure onset contacts that were stimulated with 1 Hz SPES (median stimulation intensity 4 mA, range 0-59 seconds from onset of electrical stimulation), but in none of the control or IZ_p_ contacts. Only habitual seizures and auras were recorded. Non-specific discomfort was reported in two cases. None of the evoked auras led to generalized seizures. All but one were focal with retained awareness.

### Surgical Outcomes

Stimulation in the group of control sites in multifocal or poor surgical outcome cases exhibited a higher number of evoked responses at distant sites compared to the group of localizable onsets or good surgical outcome cases (CI medians 0.5 vs. 0.12, p 0.04).

There was a statistically significant correlation between duration of epilepsy and (CI) of non-SOZ contacts up to 30 years of epilepsy (r= 0.49, p = 0.01) (Figure 2-I).

## Discussion

In this study, we demonstrated that the SOZ exhibits enhanced pathologic connectivity characterized by multi-regional or wide-spread abnormal evoked responses to single pulse electrical stimulation. This was seen both in temporal and extra-temporal epilepsies. These responses were recorded at relatively distant brain regions from the site of stimulation. This emphasizes that the increased connectivity is not a result of sampling bias. The appearance of evoked responses with stimulation of the SOZ was more complex than with control sites. The evoked responses following stimulation or recording from SOZ exhibited complex configuration and multiple peaks, which did not comply with N1-N2 of cortico-cortical evoked potentials CCEPs reported by others (see figure 2-A-F). This was further illustrated by the qualitative analysis where late and slow responses were seen almost exclusively after stimulation of epileptic brain regions. This also accounted for the statistically significant differences in latencies and duration of evoked responses between epileptic and non-epileptic brain regions, despite the little difference in absolute values. In addition, stimulation across patients differed significantly between epileptic and control sites for corresponding brain regions across different subjects, or within subjects. We successfully quantified these responses by implementing a metric which we labeled the Connectivity Index (CI), which accounts for both the spatial extent and the distance for the responses. We argue that the abnormal evoked responses observed after single pulse stimulation are a combination of a regional pathologic facilitation of synaptic transmission^8 16^, and subcortical recruitment in subjects with more wide-spread responses^2,16^.

The use of (CI) as a classifier provided a reasonable trade-off between sensitivity and specificity. Despite the inclusion of only ‘highly’ active irritative zone contacts. It is possible the differences between SOZ and IZ_p_ did not reach statistical significance due to the relatively small sample size but may indicate as well that IZ_p_ include possibly epileptogenic contacts. Outcome validated studies of the resection of subgroup of IZ_p_ and SOZ may elucidate that further. This may be further elucidated with prospectively stimulated contacts. In fact; we believe it is worthwhile to investigate the clinical utility of SPES of brain regions to localize SOZ, and to grade epileptogenicity prospectively before recording seizures (i.e. blinded probing of SOZ). Previous studies have shown that the cellular response to external electrical stimulation is enhanced in the epileptogenic cortex^13^. In vivo, cellular recordings from lateral temporal cortex in patients with intractable epilepsy showed that ‘epileptic’ neurons with spontaneous high-frequency bursts are more likely to generate evoked single unit activities after direct cortical stimulation than do ‘normal’ firing neurons^19^. Wilson et al. used evoked potentials elicited by stimulation of the limbic structures to investigate the “preferred pathway” of epileptic activity. Consistently, other reports of increased connectivity measures of SOZ as by fMRI and spontaneous electrocorticography exist^11,20^. Cortico-cortical-evoked potentials (CCEPs) employing similar technique (i.e. single pulse) often in evaluating physiologic brain connectivity typically emphasizes two physiologic early responses N1-N2 within 200 ms following stimulation. The most profound responses in our series after stimulation of epileptic brain regions were encountered > 200 ms and the pathologic responses did not respect the N1-N2 configuration, and frequently spanned the entire 1-second window hence the extended analysis time.

The correlation between the duration of epilepsy and the (CI) of non-SOZ contacts is of interest and is concordant with prior reports in mesial temporal lobe epilepsy^4^ and frontal lobe epilepsy^15^. This adds to the cumulative evidence of epilepsy being a progressive disease, at least in the majority of subset of patients undergoing icEEG for refractory epilepsy, and supports the approach of considering epilepsy surgery sooner rather than later with the caveat however that correlation does not necessarily indicate causality and there could be other clinical and/or pathology-related factors resulting in both longer intervals until surgical intervention and the degree of epileptogenicity of non-SOZ tissue. The correlation between (CI) and the stimulation intensity on the one hand, and the absence of that relationship with the distance of the site of recording of evoked responses suggests, that it is feasible, and potentially more useful to grade evoked responses from epileptic regions at even lower currents than used in this study.

The high predictive values of auras and seizures induced by electrical cortical stimulation in both temporal and extra-temporal epilepsy are comparable with and validate those reported in temporal lobe epilepsy previously^14^.

There are inherent limitations to the retrospective design of case-control studies. However, we believe this is mitigated by the prospective classification of contacts before stimulation and semi-automated analysis of evoked responses. While intracranial EEG provides a non-symmetrical and inhomogeneous sampling of brain regions, and relatively limited spatial resolution compared to other non-invasive modalities, the practice of extensive spatial sampling at our center and the use of normalized values in this study compensate partially for these limitations. It is possible that a combination of qualitative and quantitative methods of evoked responses will be more efficient in detecting and grading epileptogenicity. Employing controls as identified in our study based on the interictal profile has been successfully used in similar settings^1^. We acknowledge however, that identifying true-controls in a cohort of epilepsy patients has its own theoretical limitations in the face of modern view of epilepsy as a network disease. This holds true especially for non-lesional or subtly lesional cases.

## Conclusions

Epileptic brain regions exhibited increased connectivity, including with more distant sites, compared to non-epileptic quiet contacts. The averaged evoked responses (SPEPs) after 1 Hz stimulation of the seizure onset zone were characterized by atypical morphology, duration, and spatial distribution. Auras and seizures provoked by low frequency stimulation, though not sensitive, were highly specific to the seizure onset zone in this series. Our current study suggests these methods provide a robust supplement to current standard of care. The connectivity index provides a robust supplement to standard icEEG evaluations a resoanble tradeoff between sensitives and specifities for classification of the epileptic contacts and the findings warrant further investigations to probe the epileptic regions and validate our metric in a blinded and prospective manner in different brain sub-regions and pathologies.

## Authors’ Contribution

QY: data collection, data analysis, drafting of segments of manuscript NG: data analysis, drafting of manuscript, HPZ: data collection, drafting of the manuscript, HB: data analysis, drafting of manuscript. RA: conceptualization, design, analysis of data, drafting a significant portion of the manuscript and figures revision of the manuscript and Supervision of the study LJH: data analysis, drafting of manuscript, DDS: data analysis, drafting of manuscript, supervision of the study.

## Acknowledgements

We wish to acknowledge research support by the American Epilepsy Society (award #412064) and CTSA Grant Number KL2TR000140 from the National Center for Advancing Translational Science (NCATS), a component of the National Institute of Health (NIH) and the C.G. Swebilius Trust (all to RA). Note, the content of the manuscript is solely the responsibility of the authors and do not necessarily represent the official view of the NIH. The authors wish to thank Rebecca Khozein DOM, MS, REEG/EPT, RPSGT, RNCST, and Tamara Wing REEGT for assistance in EEG data acquisition.

## References

1. Alkawadri R, Gaspard N, Goncharova, II, Spencer DD, Gerrard JL, Zaveri H, et al: The spatial and signal characteristics of physiologic high frequency oscillations. Epilepsia 55:1986–1995, 2014

2. Arthuis M, Valton L, Regis J, Chauvel P, Wendling F, Naccache L, et al: Impaired consciousness during temporal lobe seizures is related to increased long-distance cortical-subcortical synchronization. Brain 132:2091–2101, 2009

3. Asano E, Juhasz C, Shah A, Sood S, Chugani HT: Role of subdural electrocorticography in prediction of long-term seizure outcome in epilepsy surgery. Brain 132:1038–1047, 2009

4. Bartolomei F, Chauvel P, Wendling F: Epileptogenicity of brain structures in human temporal lobe epilepsy: a quantified study from intracerebral EEG. Brain 131:1818–1830, 2008

5. Feindel W, Leblanc R, de Almeida AN: Epilepsy surgery: historical highlights 1909-2009. Epilepsia 50 Suppl 3:131–151, 2009

6. Fong JS, Alexopoulos AV, Bingaman WE, Gonzalez-Martinez J, Prayson RA: Pathologic findings associated with invasive EEG monitoring for medically intractable epilepsy. Am J Clin Pathol 138:506–510, 2012

7. Fong JS, Jehi L, Najm I, Prayson RA, Busch R, Bingaman W: Seizure outcome and its predictors after temporal lobe epilepsy surgery in patients with normal MRI. Epilepsia 52:1393–1401, 2011

8. Iwasaki M, Enatsu R, Matsumoto R, Novak E, Thankappen B, Piao Z, et al: Accentuated cortico-cortical evoked potentials in neocortical epilepsy in areas of ictal onset. Epileptic Disord 12:292–302, 2010

9. Jehi LE, O’Dwyer R, Najm I, Alexopoulos A, Bingaman W: A longitudinal study of surgical outcome and its determinants following posterior cortex epilepsy surgery. Epilepsia 50:2040–2052, 2009

10. Joshi A, Scheinost D, Okuda H, Belhachemi D, Murphy I, Staib LH, et al: Unified framework for development, deployment and robust testing of neuroimaging algorithms. Neuroinformatics 9:69–84, 2011

11. Lee HW, Arora J, Papademetris X, Tokoglu F, Negishi M, Scheinost D, et al: Altered functional connectivity in seizure onset zones revealed by fMRI intrinsic connectivity. Neurology 83:2269–2277, 2014

12. Matsumoto R, Nair DR, LaPresto E, Najm I, Bingaman W, Shibasaki H, et al: Functional connectivity in the human language system: a cortico-cortical evoked potential study. Brain 127:2316–2330, 2004

13. Meisel C, Schulze-Bonhage A, Freestone D, Cook MJ, Achermann P, Plenz D: Intrinsic excitability measures track antiepileptic drug action and uncover increasing/decreasing excitability over the wake/sleep cycle. Proc Natl Acad Sci U S A 112:14694–14699, 2015

14. Munari C, Kahane P, Tassi L, Francione S, Hoffmann D, Lo Russo G, et al: Intracerebral low frequency electrical stimulation: a new tool for the definition of the “epileptogenic area”? Acta Neurochir Suppl (Wien) 58:181–185, 1993

15. Simasathien T, Vadera S, Najm I, Gupta A, Bingaman W, Jehi L: Improved outcomes with earlier surgery for intractable frontal lobe epilepsy. Ann Neurol 73:646–654, 2013

16. Valentin A, Alarcon G, Garcia-Seoane JJ, Lacruz ME, Nayak SD, Honavar M, et al: Single-pulse electrical stimulation identifies epileptogenic frontal cortex in the human brain. Neurology 65:426–435, 2005

17. Valentin A, Alarcon G, Honavar M, Garcia Seoane JJ, Selway RP, Polkey CE, et al: Single pulse electrical stimulation for identification of structural abnormalities and prediction of seizure outcome after epilepsy surgery: a prospective study. Lancet Neurol 4:718–726, 2005

18. Vilensky JA, Gilman S: Horsley was the first to use electrical stimulation of the human cerebral cortex intraoperatively. Surg Neurol 58:425–426, 2002

19. Wyler AR, Ward AA, Jr.: Neurons in human epileptic cortex. Response to direct cortical stimulation. J Neurosurg 55:904–908, 1981

20. Zaveri HP, Pincus SM, Goncharova, II, Duckrow RB, Spencer DD, Spencer SS: Localization-related epilepsy exhibits significant connectivity away from the seizure-onset area. Neuroreport 20:891–895, 2009

